# A Reference Genome from the symbiotic hydrozoan, *Hydra viridissima*

**DOI:** 10.1101/2020.05.26.117382

**Authors:** Mayuko Hamada, Noriyuki Satoh, Konstantin Khalturin

**Author notes:** **Corresponding author:** Mayuko Hamada, Ushimado Marine Institute, Okayama University, Kashino 130-17, Ushimado, Setouchi, Okayama 701-4303, Japan.

## Abstract

Cnidarians are one of the oldest eumetazoan taxa, and are thought to be a sister group to all bilaterians. In spite of comparatively simple morphology, cnidarians exhibit diverse body forms and life histories. In addition, many cnidarian species establish symbiotic relationships with microalgae. Various *Hydra* species have been employed as model organisms since the 18th century. Introduction of transgenic and knock-down technologies made them ideal experimental systems for studying cellular and molecular mechanisms involved in regeneration, body-axis formation, senescence, symbiosis, and holobiosis. In order to provide an important reference for genetic studies, the *Hydra magnipapillata* genome was sequenced. However, the initial published version of the *H. magnipapillata* genome did not achieve assembly continuity comparable to those of other model systems, due mainly to a large number of transposable elements. For almost a decade, the highly fragmented genome assembly of *H. magnipapillata* (scaffold N50=128Kb) has remained the only genomic resource for this genus with several dozen species. Here we report a draft 280-Mbp genome assembly for *Hydra viridissima* strain A99, a symbiotic, early diverging member of the *Hydra* clade, with a scaffold N50 of 1.1 Mbp. The *H. viridissima* genome contains an estimated 21,476 protein-coding genes. Comparative analysis of Pfam domains and orthologous proteins highlights characteristic features of *H. viridissima*, such as diversification of innate immunity genes that are important for host-symbiont interactions. Thus, the *Hydra viridissima* assembly provides an important hydrozoan genome reference that will facilitate symbiosis research and better comparisons of metazoan genome architectures.

## INTRODUCTION

The Cnidaria is an evolutionarily ancient and well-defined phylum, characterized by the possession of nematocytes (Brusca et al. 2016). Cnidarian species belong to the Medusozoa, which consists of the Hydrozoa, the Scyphozoa and the Cubozoa, and the Anthozoa (Fig. 1A). Although cnidarian morphology exhibits astonishingly diverse forms and life styles, those of the fresh water Hydrozoa *Hydra* are relatively simple. It possesses only a polyp stage, while other medusozoans exhibit alternation of polyp and jellyfish generations. With its simple body structure and easy laboratory cultivation, *Hydra* has been an experimental model for studying cellular and molecular mechanisms underlying the formation of the body axis (Bode 2011), regeneration (Trembley et al. 1744; Bode 2003; Holstein et al. 2003), and also holobiotic relationships with microbiota (Deines and Bosch 2016). Introduction of transgenic and knock-down technologies further promoted these studies (Wittlieb et al. 2006). In order to provide genetic information for these studies, the brown hydra, *Hydra magnipapillata*, genome was sequenced (Chapman et al. 2010).

**Figure 1.**
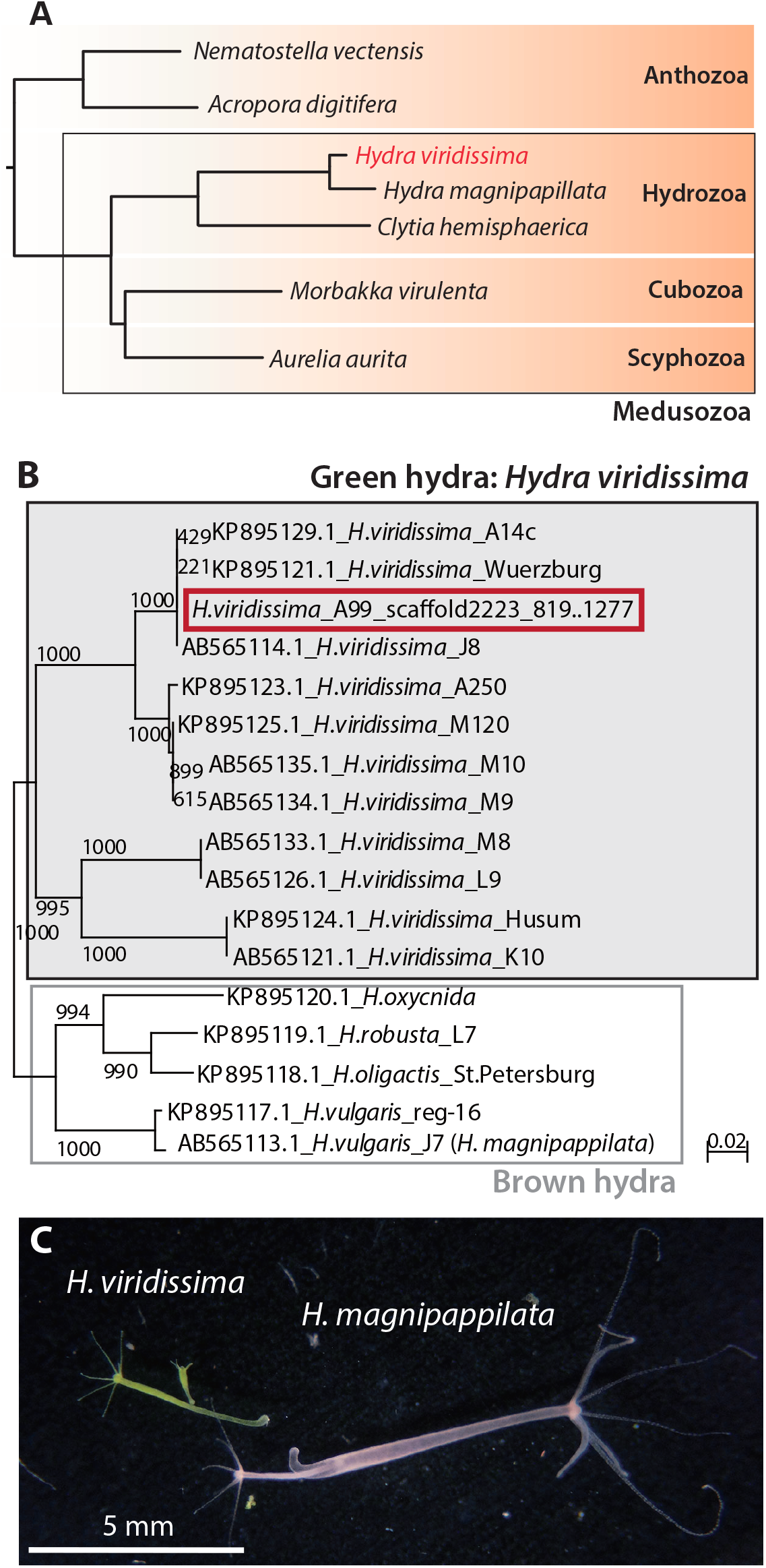
Phylogeny and morphology of green hydra *Hydra viridissima*. (A) Phylogenetic position of *H. viridissima* (red) within the phylum Cnidaria. (B) Relationship of *Hydra viridissima* strain A99 (red) with other *H. viridissima* strains and brown hydra species, based on phylogenetic analysis by NJ method using cytochrome c oxidase subunit I (COI) gene sequences. The genomic region in *H. viridissima* A99 and Genbank IDs in other strains used in the phylogenetic analysis are indicated. (c) Photograph of *H. viridissima* (left) and *H. magnipapillata* (right)*. H. viridissima* is smaller than *H. magnipapillata*, and green due to symbiotic *Chlorella* in its endodermal epithelial cells.

In addition to *H. magnipapillata*, genomes of representative species belonging to each subgroup of cnidarians have been sequenced, including the other hydrozoans *Clytia hemisphaerica* (Leclère et al. 2019), the scyphozoan jellyfish, *Aurelia aurita* (Khalturin et al. 2019; Gold et al. 2019) and the cubozoan box jellyfish, *Morbakka virulenta* (Khalturin et al. 2019), the anthozoans sea anemones *Nematostella vectensis* (Putnam et al. 2007), *Aiptasia* (Baumgarten et al. 2015), the coral *Acropora digitifera* (Shinzato et al. 2011) and other coral species. The quality of the initial published version of the *Hydra magnipapillata* genome assembly was approximately 4-22x poorer than those of other cnidarian genomes. A burst of transposable elements occurred in the brown hydra lineage, resulting in a large genome size (Chapman et al. 2010; Wong et al. 2019) and making long assembly of the genome difficult.

Compared to *H. magnipappilata*, the genome size of green hydra *Hydra viridissima* was expected to be much smaller (Zacharias et al. 2004). *H. viridissima* is the most basal species in the genus *Hydra* (Fig. 1B) (Martínez et al. 2010; Schwentner and Bosch 2015), and a symbiotic species that establishes a mutualistic relationship with microalgae to exchange metabolites (Fig. 1C) (Muscatine 1965; Cernichiari et al. 1969; Mews 1980; McAuley 1991). While symbiosis with dinoflagellates is observed in many marine cnidarians, such as corals, jellyfish, and sea anemones, *H. viridissima* harbors the green alga, *Chlorella* (Douglas and Huss 1986; Huss et al. 1993/1994; Davy et al. 2012).

While the two lineages have similar, simple body structures, as described above, there are significant differences between green and brown hydras. However, little is known about the genetics that enable green hydras to support this unique symbiosis. In addition, the evolutionary trajectory of the *Hydra* lineage among the Medusozoa is still unclear. Here we report a draft assembly of the ~284-Mbp genome of *Hydra viridissima* strain A99 as another better reference of *Hydra* genomes, and demonstrate characteristics of the green hydra genome, including transposable elements and genes involved in its body plan and in symbiosis.

## MATERIALS AND METHODS

### *Hydra* and extraction of DNA

The Australian *Hydra viridissima* strain A99, which was kindly provided by Dr. Richard Campbell, at the University of California at Irvine, was used in this study. Polyps were maintained at 18°C on a 12-hour light/dark cycle and fed with *Artemia* two or three times a week. DNA for genome sequencing were isolated from about 1000 polyps that were clonally cultured. Before genomic DNA extraction, symbiotic algae in *H. viridissima* were removed by photobleaching with 5 μM DCMU (3-(3,4-dichlorophenyl)-1,1-dimethylurea), as described previously (Pardy 1976; Habetha et al. 2003). To remove contamination from other organisms, polyps were starved and treated with antibiotics (50 mg/L ampicillin, rifampicin, neomycin, and streptomycin) for one week.

After several rounds of washing in sterilized culture medium, polyps were lysed in DNA extraction buffer (10 mM Tris-HCl, pH 8.0, 100 mM NaCl, 25 mM EDTA, pH 8.0, 0.5% SDS) and digested with 100 mg/L Proteinase K. Genomic DNA was extracted using the standard phenol-chloroform method with 100 mg/L RNaseA treatment. The quantity of DNA was determined using a NanoDrop (Thermo Fisher Scientific, Waltham, MA, USA), and the quality of high molecular-weight DNA was checked using agarose gel electrophoresis.

### Sequencing of genomic DNA

In paired-end library preparations for genome sequencing, genomic DNA was fragmented with a Focused-ultrasonicator M220 (Covaris Inc., Woburn, MA, USA). Paired-end library (average insert size: 540 bp) and mate-pair libraries (average insert sizes: 3.2, 4.6, 7.8 and 15.2 kb) were prepared using Illumina TruSeq DNA LT Sample Prep Kits and Nextera Mate Pair Sample Preparation Kits (Illumina Inc., San Diego, CA, USA), following the manufacturer’s protocols. These libraries were quantified by Real-Time PCR (Applied Biosystems StepOnePlus, Thermo Fisher Scientific) and quality controlled using capillary electrophoresis on a Bioanalyzer. Genome sequencing was performed using the Illumina Miseq system with 600-cycle chemistry (2 × 300 bp). Genome sequencing statistics is shown in Table S1A.

### RNA extraction and sequencing

Total RNA was extracted from about 1000 polyps in six different conditions (with or without symbiotic algae, in light or dark conditions, and treated with antibiotics or DMCU with symbiotic algae) using Trizol reagent (Thermo Fisher Scientific) and an RNeasy Mini kit (QIAGEN, Hilden, Germany). Quantity of RNA were checked with a NanoDrop (Thermo Fisher Scientific). Quality of total RNA was checked with a BioAnalyzer (Agilent Technologies, Santa Clara, CA, USA). For mRNA-seq, libraries were produced using an Illumina TruSeq Stranded mRNA Sample Prep Kit and were sequenced on HiSeq 2000 instruments using 2 × 150-cycle chemistry. mRNA-sequencing statistics is shown in Table S1B.

### Assembly and gene prediction

Sequencing reads of genomic DNA were assembled using Newbler Assembler version 2.8 (Roche, Penzberg, Germany) and subsequent scaffolding was performed with SSPACE (Boetzer et al. 2011). Gaps inside scaffolds were closed with paired-end and mate-pair data using GapCloser of the Short Oligonucleotide Analysis Package (Luo et al. 2012). Then one round of Haplomerger2 processing pipeline (Huang et al. 2017) was applied to eliminate redundancy in scaffolds and to merge haplotypes. For gene model prediction, we used a species-specific gene prediction model that was trained based on mapping of the *Hydra viridissima* transcriptome and raw RNAseq reads against the genome assembly. Mapping and gene structure annotation were performed using PASA pipeline v2.01 and were used to train AUGUSTUS software (Haas et al. 2003; Stanke et al. 2006). Genome completeness was evaluated using BUSCO (Benchmarking Universal Single-Copy Ortholog) (Seppey et al. 2019). RNA-Seq transcripts were mapped to the genome assembly with BWA.

### Genome size estimation

Genome size was estimated from raw paired-end reads by k-mer distribution analysis. Jellyfish v2.0.0 was used to count k-mers and their frequencies (Marcais and Kingsford 2011). The *Hydra viridissima* genome size was estimated from obtained k-mer distribution frequencies using GenomeScope web tool (Vurture et al. 2017) (http://qb.cshl.edu/genomescope/).

### Analysis of repetitive elements

Repetitive elements in the draft genome assemblies of *Hydra viridissima* were identified *de novo* with RepeatScout version 1.0.5 (http://www.repeatmasker.org/RepeatModeler) and RepeatMasker version 4.0.6 (http://www.repeatmasker.org). Repetitive elements were filtered by length and occurrence so that only sequences longer than 50 bp and present more than 10 times in the genome were retained. The resulting sets of repetitive elements were annotated using BLASTN and BLASTX searches against RepeatMasker.lib (35,996 nucleotide sequences) and RepeatPeps.lib (10,544 peptides) bundled with RepeatMasker version 4.0.6. The results of both searches were combined, and BLASTX results were given priority in cases where BLASTN and BLASTX searches gave conflicting results.

### Analysis of *Hydra viridissima* genes

For comparative analysis of *H. viridissima* genes among cnidarians, protein sequences were obtained from Compagen (http://www.compagen.org/) for *Hydra magnipapillata* (Chapman et al. 2010), from JGI (https://genome.jgi.doe.gov/Nemve1/Nemve1.home.html) for *Nematostella vectensis* (Putnam et al. 2007), from MARIMBA (available at http://marimba.obs-vlfr.fr/organism/Clytia/hemisphaerica) for *Clytia hemisphaerica* (Leclère et al. 2019), from the genome project website of OIST Marine Genomics Unit (https://marinegenomics.oist.jp/gallery/gallery/index) for *Acropora digitifera* (Shinzato et al. 2011), and for *Morbakka virulenta* and *Aurelia aurita* in Atlantic Ocean (Khalturin et al. 2019). In comparative analyses, domain searches against the Pfam database (Pfam-A.hmm) were performed using HMMER (Finn et al. 2016), and ortholog gene grouping was done using OrthoFinder (Emms and Kelly 2015). To classify homeodomain containing proteins, BLAST searches and phylogenetic analysis were performed. Homeodomain sequences in various animals were obtained from Homeobox Database (http://homeodb.zoo.ox.ac.uk/families.get?og=All) were used (Zhong and Holland 2011).

For phylogenetic analysis, multiple alignments were produced with CLUSTALX (2.1) with gap trimming (Larkin et al. 2007). Sequences of poor quality that did not align well were deleted using BioEdit (Hall 1999) Phylogenetic analyses were performed using the Neighbor-Joining method (Saitou and Nei 1987) in CLUSTALX with default parameters (1,000 bootstrap tests and 111 seeds). Representative phylogenetic trees were drawn using FigTree v1.4.4 (http://tree.bio.ed.ac.uk/software/figtree/). Gene/protein ID used for phylogenetic analysis are shown in the trees (Figs S2 and S3).

### Data availability

This whole-genome shotgun sequencing project has been deposited at DDBJ/ENA/GenBank under the accession QPEY00000000 (BioProject ID: SAMN09635813). RNA-seq reads has been deposited at SRA of NCBI (SRX6792700-SRX6792705). Genome sequences, gene models, and a genome browser are also accessible at the website of the OIST Marine Genomics Unit Genome Project (https://marinegenomics.oist.jp/hydraviridissima_A99). The genome browser was established for assembled sequences using the JBrowser 1.12.3 (Skinner et al. 2009).

Gene annotations from the protein domain search and BLAST search are likewise shown on the site. Reagents, software and datasets used in this study were listed in Reagent Table. Supplemental files are available at FigShare; Figure S1. k-mer frequency distribution plots in *Hydra viridissima* A99 genome sequence, Figure S2. Phylogenetic analysis of ANTP genes, Figure S3. Phylogenetic analysis of PRD genes, Table S1. Sequencing statistics for *Hydra viridissima* A99, Table S2. Summary of repetitive sequences in the *Hydra viridissima* A99 genome assembly, Table S3. Number of Pfam domain-containing genes in the *Hydra viridissima* A99 genome, Table S4. Number of orthologs enriched in *Hydra viridissima* A99 (A) and *Hydra* (B), Table S5. Gene IDs of ANTP genes in *Hydra viridissima* A99.

## RESULTS AND DISCUSSION

### Genome architecture of *Hydra viridissima*

*Hydra viridissima* appears green because of the symbiotic *Chlorella*, and has a smaller body compared to the brown hydra *Hydra magnipapillata* (Fig. 1C). We decoded the genome of *H. viridissima* strain A99, which is closely related to strain A14c, Wuerzburg and J8 (Fig. 1B). We previously reported the genome of its specific symbiotic algae, *Chlorella* sp. A99, and demonstrated the metabolic co-dependency between *H. viridissima* A99 and the symbiont (Hamada et al. 2018).

The genome of *H. viridissima* A99 was sequenced using the Illumina platform with paired-end and mate pair libraries. Statistics of sequence reads, the assembly, and genome architecture are shown in Table 1. We obtained ~7,070 Mbp of paired-end sequences, and 4,765, 4,769, 3,669 and 3,551 Mbp for 3.2k, 4.6k, 7.8k, and 15.2k insert-size pate-pair sequences, respectively, comprising a total of ~23,826 Mbp (Table S1). The size of the *H. viridissima* genome was estimated at ~254 Mbp using k-mer analysis (k-mer=19) based on paired-end sequence data (Fig. S1). This indicates that we achieved more than 90-fold sequence coverage of the genome. On the other hand, the total length of the genome sequence assembly reached 284,265,305 bp. That is, the total assembly represented a good match for the estimated genome size.

**Table 1.** Comparison of the genome assembly statistics of cridarians.

|  | Hydrozoa |  |  | Scyphozoa | Cubozoa | Anthozoa |  |
| --- | --- | --- | --- | --- | --- | --- | --- |
| Species | *Hydra viridissima* *1 | *Hydra magnipapillata* *2 | *Clytia hemisphaerica* *3 | *Aurelia aurita* *4 | *Morbakka virulenta* *4 | *Acropora digitifera* *5 | *Nematostella vectensis* *6 |
| Geographical origin | Laboratory strain (A99) | Laboratory strain | Villefranche (Atlantic) | Baltic Sea (Atlantic) | Seto Inland Sea (Pacific) | Okinawa (Pacific) | Laboratory strain |
| Genome size (Mbp) | 284 | 852 | 445 | 377 | 952 | 420 | 457 |
| Number of Scaffolds | 2,677 | 20,914 | 7,644 | 2,710 | 4,538 | 2,420 | 10,804 |
| Longest scaffold (Mbp) | 5.1 | 0.91 | 2.9 | 4.4 | 14.5 | 2.5 | 3.3 |
| Scaffold N50 (Mbp) | 1.1 | 0.1 | 0.4 | 1.0 | 2.2 | 0.5 | 0.5 |
| GC content (%) | 24.73 | 25.41 | 35 | 34.65 | 31.37 | 39 | 39 |
| Repeats (%) | 37.5 | 57 | 41 | 44.67 | 37.41 | 12.9 | 26 |
| Gap rate (%) | 16.77 | 7.82 | 16.6 | 6.63 | 11.87 | 15.2 | 16.6 |
| Number of genes | 21,476 | 31,452 | 26,727 | 28,625 | 24,278 | 23,668 | 27,273 |
| Mean gene length (bp) | 7,637 | 6,873 | 5,848 | 10,215 | 21,444 | 8,727 | 4,500 |
| Mean exon length (bp) | 209 | 247 | 281 | 368 | 350 | 316 | 208 |
| Mean intron length (bp) | 838 | N/A | N/A | 1391 | 3572 | 1057 | 800 |
| BUSCO (complete) % | 83.9 | 76.8 | 86.4 | 79.8 | 81.5 | 74.5 | 91.4 |

\*1, This study. \*2, Chapman et al., 2010. \*3, Leclerc et al., 2019. \*4, Khaltunin et al., 2019. \*5, Shinzato et al., 2011. \*6, Putnam et al., 2007.

Although genomic DNA was extracted from a clonally propagated culture of hydra polyps maintained in the laboratory, heterozygosity was comparatively high (2.28% of the entire sequence) (Fig. S1). Thus, polyps originally collected from the wild had a high level of heterozygosity. Repetitive sequences constituted 37.5 % of the genome and the gap rate was 16.8% of the genome (Table 1; see next section). Scaffolds from the present analysis numbered 2,677 and the scaffold N50 was 1.1 Mbp, with the longest scaffold reaching 5.1 Mbp (Table 1). The GC content of the genome was 24.7% (Table 1), suggesting that *H. viridissima* has an AT-rich genome. Using 67,339,858,036 nucleotides of RNA-sequence data (Table S1), we predicted gene models. The genome was estimated to contain 21,476 protein-coding genes (Table 1). We did not find any gene models with sequence similarities to the symbiotic *Chlorella*. The mean gene length, exon length, and intron length were 7637, 209, and 838 bp, respectively (Table 1). The BUSCO method supports 84% of the *H. viridissima* gene models as complete and predicts that the genome accounts for 91% of the genes, including partial sequences (Table 1).

During assembly and gene annotation, we noticed that scaffold2223, composed of 18,375 base pairs (bp), contained almost the entire *H. viridissima* mitochondrial genome. The mitochondrial genome of *H. viridissima* strain A99 was linear, as reported by Pan et al. (2014) for the other green hydra *Hydra sinensis*, while the brown hydra, *Hydra vulgaris*, mitochondrial genome is composed of two linear molecules (Pan et al. 2014a; Pan et al. 2014b).

Comparison of *H. viridissima* genome statistics with those of other cnidarian genomes showed that the *H. viridissima* genome assembly is of higher quality than the *H. magnipapillata* genome assembly (Table 1). Although the genome size of *H. magnipapillata* is approximately 3x larger than that of *H. viridissima*, the number of scaffolds and the scaffold N50 were ten-times better in *H. viridissima* (2,677 and 1.1 Mbp, respectively) than in *H. magnipapillata* (20,914 and 0.1 Mbp, respectively) (Table 1). BUSCO support for gene modeling was higher in *H. viridissima* (83.9%) than in *H magnipapillata* (76.8%) (Table 1). Compared to *H. magnipapillata*, the green hydra has a compact genome, with ~65% as many genes (Table 1).

### Repetitive sequences in the *Hydra viridissima* genome

Although the abundance of repetitive sequences in anthozoan genomes is generally low (15~17%), genomes of medusozoans and hydrozoans have comparatively high levels of repetitive sequences, 57% in *H. magnipapillata*, 41% in *Clytia*, 45% in *Aurelia*, and 37% in *Morbakka* (Table 1). This was also true of *H. viridissima* (37.5%) (Table 1). DNA transposons were the most abundant type, accounting for approximately 22.41% of the genome (Figure 2A, Table S2). Of these, TcMariner, CMC, Maverick and hAT were largest components (Figure 2A). On the other hand, the percentages of LTR retrotransposons (1.63%) and non-LTR retrotransposons (0.99%) were comparatively low (Figure 2A, Table S2).

**Figure 2.**
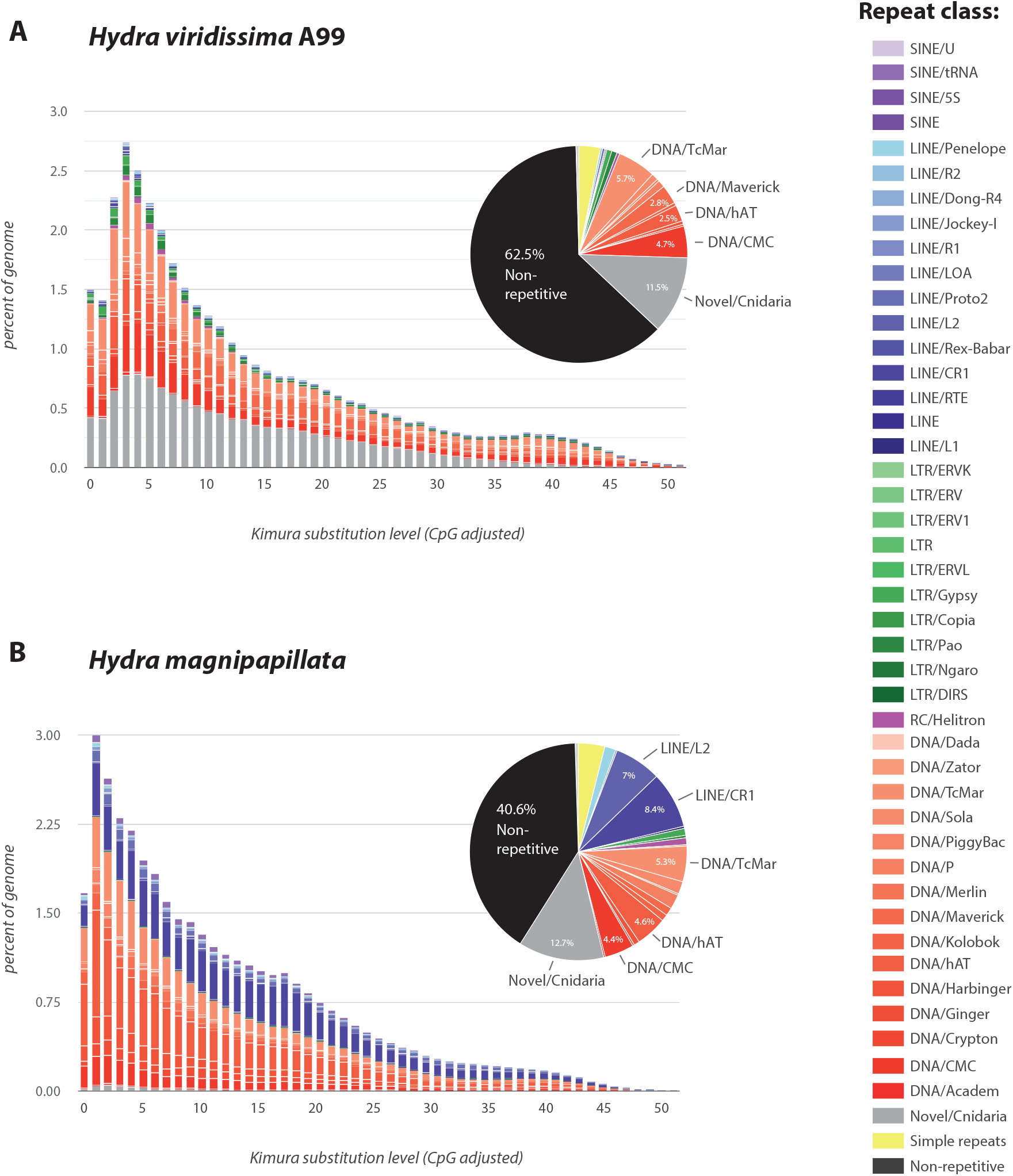
Interspersed Repeat Landscape in *Hydra*. Components and proportions of repetitive sequences in the genome of (A) *Hydra viridissima* A99 and (B) *H. magnipapillata* are shown. Classes of repeat are shown in the right column.

In comparing repetitive elements between *H. viridissima* and *H. magnipapillata*, it first becomes apparent that DNA transposons (DNA/TcMar, DNA/hAT and DNA/CMC) occupy a similar portion of both *Hydra* genomes (Figure 2). In addition, novel and potentially cnidarian-specific repetitive elements occupy ~12% of both genomes. Second, long, interspersed nuclear elements LINE/L2 (~7%) and LINE/CR1 (8.4%) are large components of the *H. magnipapillata* genome, but they are almost absent in the *H. viridissima* genome. It was suggested that a burst of retrotransposons occurred in the brown hydra lineage and may have caused the large genome size (Chapman et al. 2010; Wong et al. 2019). It is likely that the *H. viridissima* genome represents something close to the ancestral state of the *Hydra* genome. Molecular and evolutionary mechanisms involved in the insertion of LINE components in the *H. magnipapillata* genome will be a subject of future studies in relation to diversification and speciation within the *Hydra* clade.

### Innate immunity-related protein genes in the *Hydra viridissima* genome

Using the Pfam-domain search method, we surveyed genes for protein domains in the *H. viridissima* genome. We found approximately 4,500 different Pfam domains in this species (Table S3), a number comparable to those of other cnidarians. With the criterion that the number of domains should be ≥2x higher in the green hydra genome than those of non-symbiotic cnidarians and that the number represents enrichment by Chi-Square test (p-value<0.001), we searched for Pfam domains that are enriched in the *H. viridissima* genome (Table 2A). Then we checked the number of *H. viridissima*-enriched domains in the genome of the coral, *Acropora digitifera*, since it is also a symbiotic cnidarian. Death and NACHT were the two most highly enriched domains and TIR occurred in the top 10 (Table 2A). Death, NACHT and TIR domains are involved in the innate immune system. NACHT and TIR domains are found in the pattern-recognition receptors, Nod-like and Toll-like, which are sensors for pathogen and damage-associated molecules. The Death domain occurs in some pattern recognition receptors and their adaptors, which mediate signals to downstream immune responses. It appears that these domains are also enriched in *Acropora digitifera*, but not in non-symbiotic cnidarian species (Table 2A).

**Table 2.** Number of genes with Pfam domains enriched in the *Hydra viridissima* genome and comparison of their number in the other cnidarian genomes.

A. Domains enriched in *Hydra viridissima*
| Domain | Hv | Hm | Ch | Aa | Mv | Nv | Ad | Chi test* |
| --- | --- | --- | --- | --- | --- | --- | --- | --- |
| Death | 198 | 71 | 14 | 20 | 16 | 20 | 132 | 5.E-111 |
| NACHT | 161 | 38 | 42 | 45 | 23 | 39 | 458 | 2.E-71 |
| PIF1 | 121 | 25 | 59 | 49 | 3 | 8 | 18 | 3.E-65 |
| ATPase_2 | 64 | 20 | 28 | 14 | 13 | 19 | 36 | 1.E-19 |
| PAX | 61 | 23 | 3 | 4 | 23 | 9 | 8 | 6.E-35 |
| TIR_2 | 49 | 10 | 19 | 24 | 15 | 17 | 49 | 1.E-11 |
| Endonuclease_7 | 46 | 5 | 3 | 1 | 5 | 1 | 0 | 1.E-48 |
| RAG1 | 42 | 4 | 1 | 4 | 0 | 1 | 0 | 8.E-49 |
| TIR | 41 | 6 | 3 | 15 | 11 | 12 | 36 | 4.E-15 |
| RVT_3 | 33 | 1 | 7 | 10 | 14 | 1 | 1 | 2.E-20 |
| CbiA | 23 | 6 | 6 | 7 | 6 | 6 | 2 | 2.E-08 |
| DUF2738 | 21 | 0 | 1 | 0 | 0 | 1 | 0 | 4.E-28 |
| MarR_2 | 21 | 5 | 4 | 0 | 0 | 4 | 3 | 6.E-15 |
| HTH_7 | 21 | 7 | 2 | 4 | 0 | 1 | 1 | 2.E-15 |
| Sigma70_r4 | 17 | 5 | 1 | 1 | 1 | 0 | 2 | 6.E-14 |
| HTH_Tnp_IS630 | 15 | 4 | 1 | 1 | 1 | 1 | 0 | 3.E-12 |
| HTH_3 | 13 | 3 | 1 | 2 | 5 | 2 | 1 | 9.E-07 |
| TMEM151 | 10 | 4 | 2 | 3 | 0 | 3 | 6 | 2.E-03 |
| CIDE-N | 10 | 2 | 1 | 3 | 2 | 2 | 1 | 9.E-05 |
| DUF4807 | 7 | 0 | 1 | 3 | 1 | 3 | 7 | 1.E-02 |
| DUF1294 | 5 | 1 | 1 | 2 | 1 | 0 | 0 | 1.E-02 |
| A_deaminase | 9 | 2 | 4 | 3 | 3 | 3 | 5 | 3.E-02 |
| HD_4 | 4 | 1 | 1 | 1 | 1 | 0 | 0 | 4.E-02 |

B. Domains enriched in *Hydra* species
| Domain | Hv | Hm | Ch | Aa | Mv | Nv | Ad | Chi test* |
| --- | --- | --- | --- | --- | --- | --- | --- | --- |
| DDE_3 | 365 | 462 | 16 | 16 | 31 | 6 | 17 | 0.E+00 |
| Dimer_Tnp_hAT | 296 | 549 | 11 | 52 | 44 | 21 | 27 | 0.E+00 |
| HTH_Tnp_Tc3_2 | 266 | 279 | 15 | 13 | 23 | 0 | 3 | 1.E-251 |
| HTH_32 | 185 | 104 | 14 | 4 | 25 | 4 | 9 | 4.E-149 |
| ANAPC3 | 182 | 166 | 61 | 36 | 37 | 46 | 44 | 1.E-68 |
| HTH_23 | 166 | 137 | 6 | 15 | 33 | 8 | 20 | 1.E-114 |
| zf-BED | 116 | 83 | 14 | 9 | 8 | 19 | 5 | 3.E-78 |
| HTH_29 | 95 | 81 | 7 | 1 | 19 | 2 | 5 | 1.E-73 |
| HTH_psq | 93 | 105 | 3 | 6 | 3 | 1 | 1 | 3.E-92 |
| HTH_28 | 85 | 54 | 2 | 5 | 6 | 6 | 11 | 3.E-63 |
| SRP_TPR_like | 83 | 93 | 6 | 1 | 2 | 1 | 1 | 4.E-83 |
| DUF4806 | 69 | 57 | 6 | 1 | 0 | 0 | 0 | 2.E-65 |
| BTAD | 32 | 37 | 7 | 3 | 1 | 3 | 3 | 2.E-22 |
| Sigma70_r4_2 | 28 | 24 | 2 | 2 | 0 | 1 | 3 | 2.E-21 |
| HTH_Tnp_Tc3_1 | 26 | 26 | 1 | 0 | 0 | 0 | 0 | 1.E-25 |
| CD225 | 21 | 21 | 9 | 4 | 2 | 7 | 7 | 1.E-06 |
| IGFBP | 15 | 23 | 7 | 4 | 5 | 1 | 3 | 4.E-07 |
| Sulfate_transp | 15 | 21 | 2 | 5 | 4 | 4 | 5 | 4.E-06 |
| DUF2961 | 14 | 49 | 2 | 3 | 3 | 1 | 3 | 4.E-27 |
| DUF1280 | 13 | 20 | 4 | 4 | 0 | 3 | 2 | 1.E-07 |
| Polysacc_deac_1 | 13 | 14 | 3 | 1 | 6 | 4 | 2 | 8.E-05 |
| DUF4817 | 12 | 20 | 2 | 1 | 0 | 0 | 0 | 1.E-12 |
| GRDP-like | 9 | 7 | 2 | 1 | 2 | 2 | 3 | 7.E-03 |
| Transposase_mut | 8 | 13 | 1 | 3 | 1 | 0 | 0 | 3.E-06 |
| CARDB | 5 | 7 | 1 | 0 | 0 | 1 | 2 | 6.E-03 |
| SecA_DEAD | 3 | 9 | 1 | 1 | 1 | 0 | 0 | 8.E-04 |

Hv: *Hydra viridissima* A99, Hm: *Hydra magnipapillata*, Ch: *Clytia hemisphaerica*, Mv: *Morbakka virulenta*, Aa: *Aurelia aurita*, Nv, *Nematostella vectensis*, Ad: *Acropora digitifera*, \*Chi-test: evaluate of Chi-square test

Expansion of genes for NACHT-containing proteins is supported by identification of orthologous protein groups by OrthoFinder. Genes for proteins similar to Nod-like receptor were the second most overrepresented group in the *H. viridissima* genome (Table 3A, Table S4). However, these orthologs were not scored in the *Acropora* genome, suggesting that the NACHT-containing proteins expanded in *H. viridissima* are different from those expanded in *Acropora*.

**Table 3.**
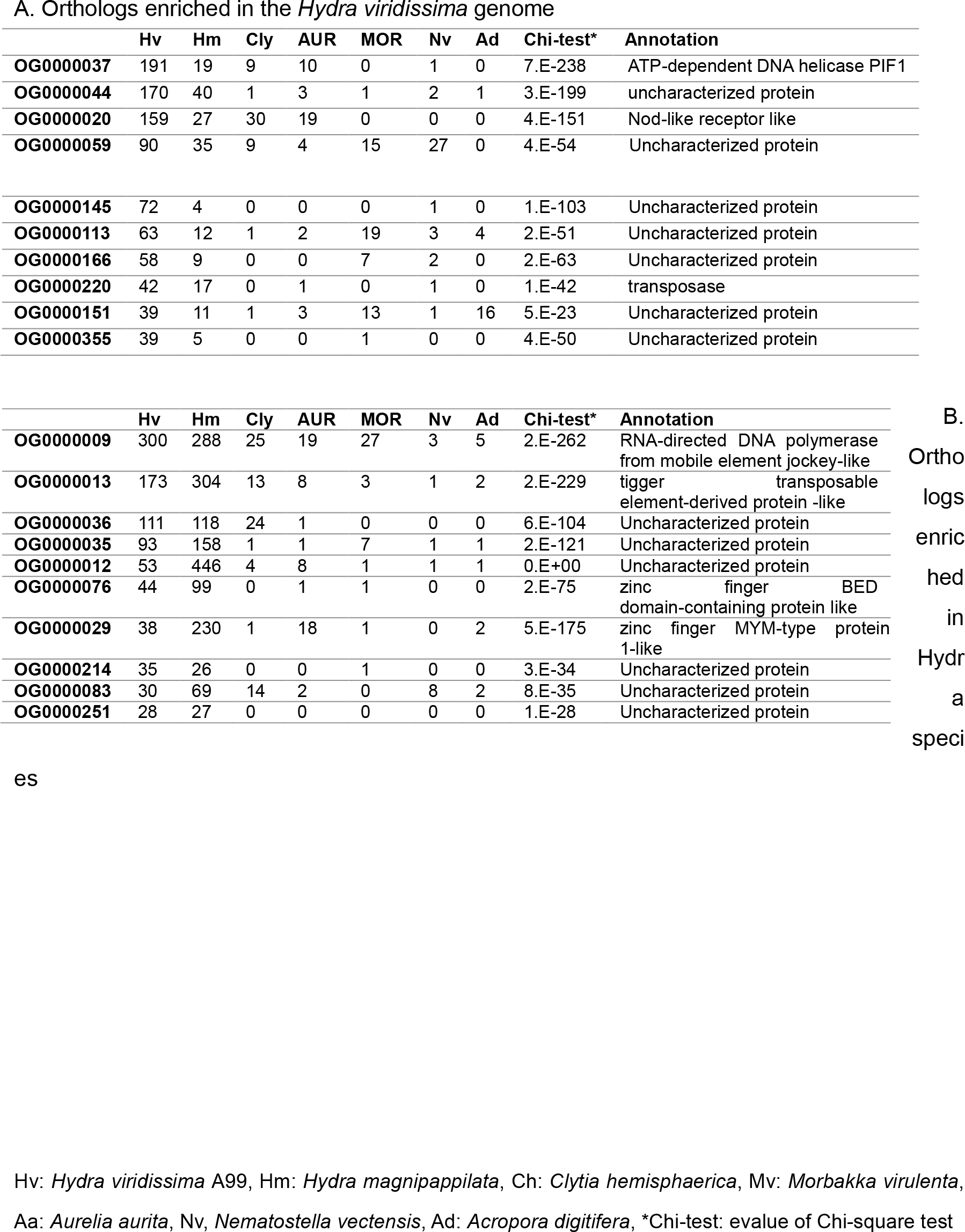
Top 10 overrepresented orthologs in the *Hydra viridissima* genome and comparison of their gene number in the other cnidarian genomes.

In *Acropora*, we previously showed unique, complex domain structures of proteins with NACHT or NB-ARC domains, which have similar structures and functions to NACHT (Hamada et al. 2013). Thus, we further examined domain combinations of NACHT/NB-ARC proteins in *H. viridissima* to determine whether such complex domain structures are also found in *H. viridissima*. Basically Nod-like receptors have a tripartite domain structure, consisting of effector-binding domain constituted of apoptosis-related domains, such as Death and DED in the N-terminus, NACHT/NB-ARC in the center, and a repeat domain that recognizes pathogen and damage-associate molecules at the C-terminus. Humans have approximately 20 Nod-like receptor family proteins, and their ligand recognition region is a leucine-rich repeat (LRR). On the other hand, in Nod-like receptors of basal metazoans, not only LRR, but also tetratricopeptide repeats (TPR), WD40 repeats, and ankyrin repeats (Ank) are found as repeat domains. We previously showed that *Acropora* has all 4 types of Nod-like receptors, and that with LRR type is the most common (Table. 4) (Hamada et al. 2013). In other cnidarians examined, only TPR and WD40 are found as repeat domains of Nod-like receptors, suggesting loss of the other types. Especially in *H. viridissima*, a larger number of genes for Nod-like receptors with TPR were found. In addition, their domain structures in *H. viridissima* vary widely, compared to those of *H. magnipapillata* (Fig. 3). For example, the domain combination of NACHT/NB-ARC with DED or TIR, was found in *H. viridissima*, but not in *H. magnipapillata*. In addition to NACHT-containing protein, *H. viridissima* has more genes for TIR domain-containing proteins, including an interleukin-1 receptor (ILR), which are not found in *H. magnipapillata*.

**Figure 3.**
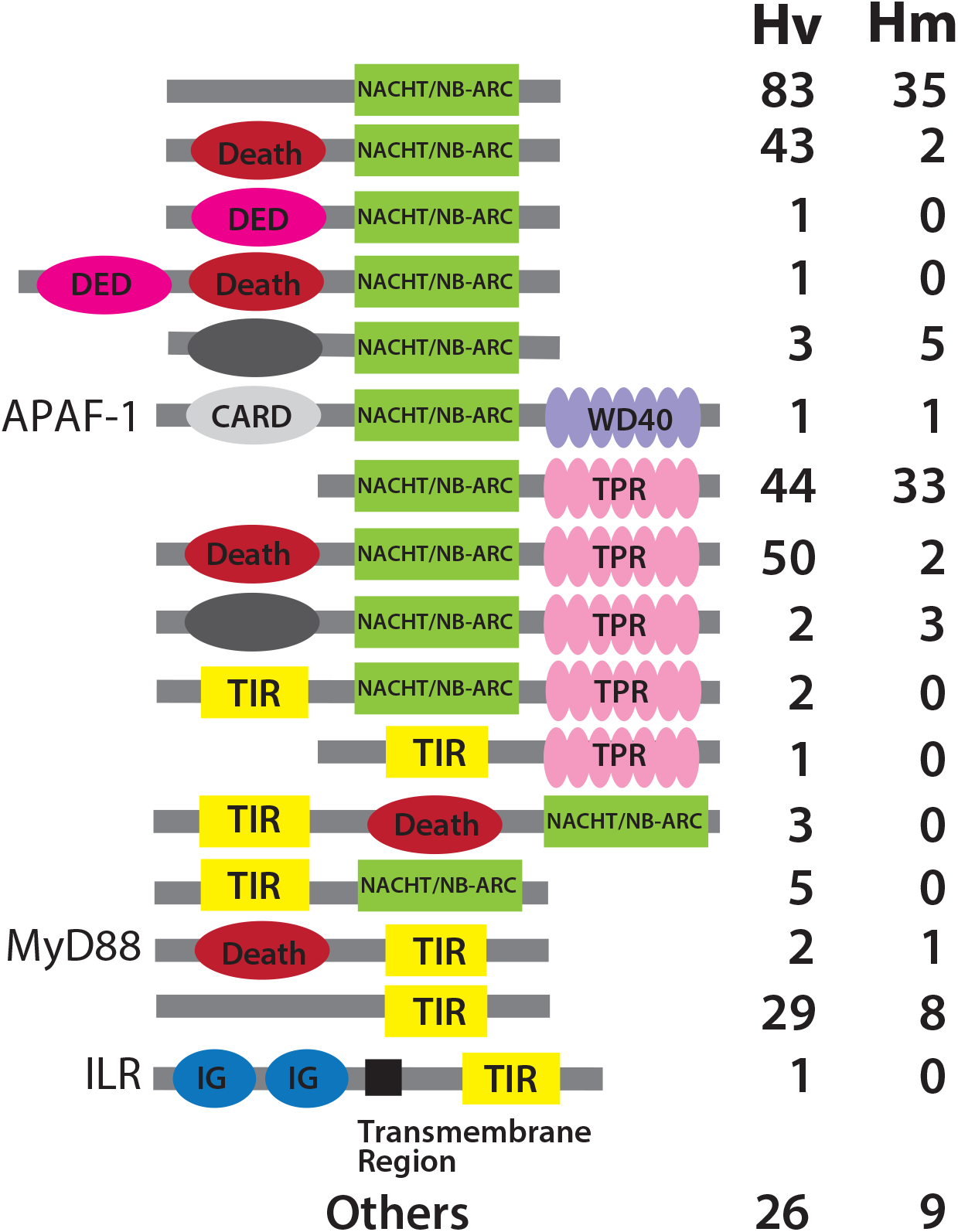
Schematic representation of domain structures of NACHT/NB-ARC or TIR-domain-containing proteins identified in *Hydra.* The domain structures and the number of NACHT/NB-ARC or TIR-domain-containing proteins in *Hydra viridissima* A99 (Hv) and *H. magnipapillata* (Hm) are shown.

**Table 4.** The number of NACHT/NB-ARC domain-containing proteins in cnidarians and the combination of repeat domains.

|  | Hv | Hm | Ch | Aa | Mv | Nv | Ad |
| --- | --- | --- | --- | --- | --- | --- | --- |
| **Total** | 264 | 89 | 56 | 58 | 24 | 6 | 489 |
| **TPR** | 103 | 38 | 3 | 8 | 4 | 2 | 57 |
| **WD40** | 2 | 4 | 3 | 2 | 3 | 1 | 8 |
| **LRR** | 0 | 0 | 0 | 0 | 0 | 0 | 125 |
| **Ank** | 0 | 0 | 0 | 0 | 0 | 0 | 11 |

Hv: *Hydra viridissima* A99, Hm: *Hydra magnipapillata*, Ch: *Clytia hemisphaerica*, Mv: *Morbakka virulenta*, Aa: *Aurelia aurita*, Nv, *Nematostella vectensis*, Ad: *Acropora digitifera*,

As mentioned above, diversification of pattern-recognition receptor-related genes are found in both *H. viridissima* and *Acropora digitifera*. Their most significant shared attribute is symbiosis, the former with *Chlorella* and the latter with the dinoflagellate, *Symbiodinium.* Therefore, it is likely that the evolutionary acquisition of symbiosis by certain cnidarians requires expansion and greater sophistication of innate immunity genes. They may participate in recognition and maintenance of symbiotic organisms in cnidarian tissues. On the other hand, the structures (e.g. repeat combination) of the Nod-like receptors most abundant in green hydra and coral were different. This indicates that species-specific adaptations to the environment occurred in each of these lineages independently.

### Genes enriched in the genus *Hydra*

We further examined Pfam domains overrepresented specifically in *H. viridissima* and others present in both *H. viridissima* and *H. magnipapillata*. This was done using the same criterion as above, that is, that the number of domains is ≥2x higher than those in other cnidarians and that the number is enriched by Chi-Square test (p-value<0.001). (Table 2B).

Pfam domain searches and ortholog protein grouping demonstrated that *H. viridissima and H. magnipapillata* possess many genes encoding domains that function in DNA binding. For example, genes containing transposase-related domain (DDE_3, Dimer_Tnp_hAT and Transposase_mut) and DNA-binding motif (HTH: helix-turn-helix and zf: zinc finger) were overrepresented in both *H. viridissima* and *H. magnipapillata* (Table 2B). In addition, PAX, Endonuclease_7, RVT_3, MarR_2, some HTHs, Sigma70_r4, CIDE-N, PIF-1 and RAG1, which bind DNA, were specifically overrepresented in *H. viridissima* (Table 2A). Particularly, ortholog protein grouping suggested that the most overrepresented gene in the *H. viridissima* genome was PIF1 ATP-dependent DNA helicase-like genes (Table 3A), which are required for maintenance of genome structure. Although the functions of these genes are presently unknown, they may be involved in the genome structure of *Hydra*, which contains many transposable elements.

Pfam domain searches also demonstrated that genes for proteins containing Sulfate_transp domain and those containing Polysacc_deac_1 domain were enriched in both *H. viridissima* and *H. magnipappilata* (Table 2B). Sulfate_transp is found in the sulfate permeases family, which is involved in uptake or exchange of inorganic anions, such as sulfate. So far, their functions in *Hydra* are unknown, but they may relate to their limnetic life style, which requires active ion uptake. Polysacc_deac_1 is found in polysaccharide deacetylase, including chitin deacetylase, which is involved in chitin metabolism. It may contribute to construction of the extracellular matrix surrounding the body or structure of nematocyte, or molecular recognition events such as immune response to pathogens with chitinous cell wall (Balasubramanian et al. 2012; Elieh Ali Komi et al. 2018; Rodrigues et al. 2016).

### Gene families for transcription factors and signaling molecules

Using Pfam-supported families, we examined the number of gene families for putative transcription regulator genes and signaling molecules (Table 5), since these genes are essential in development and physiology of metazoans. While major signaling pathways are present in cnidarians, some specialization in Cnidaria is known. For example, Wnt genes, which are important for oral-aboral body axis formation, diversified in the cnidarian lineage (Kusserow et al. 2005; Khalturin et al. 2019). Table 5A shows numbers of putative transcription factor genes in the *H. viridissima* genome. Zinc finger proteins (C2H2 type) were most abundant, with 105 members, although the abundance of this family has been noted in other cnidarian genomes (Khalturin et al. 2019). There were 33 HLH domain-containing and 50 homeobox domain-containing genes (Homeodomain-containing genes of *H. viridissima* are discussed in the next section). A similar analysis of putative signaling molecule genes showed that the *H. viridissima* genome contains 16 fibroblast growth factor (FGF)-like domain genes, 11 transforming growth factor-beta (TGB-b) genes, and 10 Wnt genes (Table 5B). These numbers are comparable to those in *H.magnipappilata*. In general, the number of transcriptional factor and signal molecule family members appeared similar among cnidarians, although a few families such as AT_hook and Hairly-orange of transcriptional factors (Table 5A) and Interleukin 3 (IL3) families of signal molecules (Table 5B) were not found in *Hydra* genomes.

**Table 5.** Number of putative transcriptional factor genes (A) and signal molecule genes (B) in the *Hydra viridissima* genome.

| Domain | Hv | Hm | Ch | Aa | Mv | Nv | Ad |
| --- | --- | --- | --- | --- | --- | --- | --- |
| ARID | 8 | 7 | 10 | 10 | 8 | 5 | 8 |
| AT_hook | 0 | 0 | 1 | 2 | 0 | 0 | 0 |
| bZIP_1 | 26 | 26 | 26 | 22 | 25 | 36 | 29 |
| bZIP_2 | 25 | 22 | 31 | 24 | 27 | 32 | 17 |
| CUT | 1 | 0 | 3 | 1 | 1 | 2 | 1 |
| DM | 6 | 6 | 7 | 9 | 11 | 12 | 7 |
| Ets | 9 | 11 | 13 | 21 | 14 | 16 | 12 |
| Forkhead | 17 | 17 | 19 | 15 | 26 | 34 | 22 |
| GATA | 4 | 4 | 3 | 7 | 7 | 4 | 5 |
| Hairy_orange | 0 | 0 | 1 | 4 | 2 | 6 | 7 |
| HLH | 33 | 36 | 44 | 52 | 50 | 72 | 53 |
| HMG_box | 30 | 33 | 31 | 30 | 29 | 33 | 27 |
| Homeobox | 50 | 44 | 70 | 88 | 82 | 153 | 96 |
| Hormone_recep | 9 | 9 | 12 | 12 | 9 | 20 | 9 |
| P53 | 2 | 1 | 2 | 1 | 2 | 3 | 3 |
| PAX | 61 | 23 | 3 | 4 | 23 | 9 | 8 |
| Pou | 2 | 3 | 3 | 4 | 4 | 6 | 4 |
| RHD_DNA_bind | 2 | 1 | 5 | 2 | 2 | 3 | 2 |
| SRF-TF | 2 | 2 | 4 | 3 | 2 | 3 | 1 |
| T-box | 6 | 7 | 11 | 10 | 9 | 14 | 10 |
| TF_AP-2 | 1 | 1 | 2 | 3 | 2 | 1 | 2 |
| zf-C2H2 | 105 | 123 | 244 | 233 | 118 | 169 | 90 |
| zf-C2HC | 1 | 2 | 3 | 2 | 3 | 3 | 4 |
| zf-C4 | 9 | 8 | 11 | 8 | 9 | 19 | 12 |

B. Signal molecule genes
| Domain | Hv | Hm | Ch | Aa | Mv | Nv | Ad |
| --- | --- | --- | --- | --- | --- | --- | --- |
| Cbl_N | 1 | 1 | 2 | 0 | 1 | 0 | 1 |
| Cbl_N2 | 1 | 1 | 1 | 0 | 1 | 0 | 1 |
| Cbl_N3 | 1 | 1 | 2 | 2 | 1 | 0 | 1 |
| DIX | 1 | 1 | 3 | 3 | 3 | 4 | 2 |
| FGF | 16 | 12 | 10 | 18 | 13 | 13 | 13 |
| Focal_AT | 1 | 2 | 1 | 1 | 1 | 0 | 0 |
| G-alpha | 29 | 28 | 29 | 29 | 32 | 37 | 22 |
| G-gamma | 2 | 2 | 1 | 7 | 5 | 4 | 3 |
| IL3 | 0 | 0 | 1 | 0 | 0 | 0 | 0 |
| PDGF | 1 | 2 | 5 | 2 | 3 | 1 | 6 |
| Phe_ZIP | 1 | 0 | 1 | 1 | 1 | 0 | 1 |
| Rabaptin | 1 | 2 | 1 | 1 | 1 | 1 | 1 |
| RGS | 13 | 13 | 14 | 16 | 16 | 13 | 11 |
| RGS-like | 2 | 3 | 1 | 2 | 1 | 0 | 1 |
| STAT_alpha | 1 | 0 | 1 | 1 | 2 | 2 | 1 |
| STAT_bind | 2 | 1 | 1 | 1 | 1 | 1 | 1 |
| STAT_int | 2 | 1 | 1 | 0 | 1 | 0 | 1 |
| TGF_beta | 11 | 11 | 7 | 9 | 9 | 7 | 10 |
| TGFb_propeptide | 8 | 10 | 4 | 6 | 8 | 6 | 9 |
| wnt | 10 | 13 | 12 | 15 | 17 | 26 | 15 |

Hv: *Hydra viridissima* A99, Hm: *Hydra magnipapillata*, Ch: *Clytia hemisphaerica*, Mv: *Morbakka virulenta*, Aa: *Aurelia aurita*, Nv, *Nematostella vectensis*, Ad: *Acropora digitifera*

### Hox and Para-Hox genes in *Hydra viridissima*

Among transcriptional factors, homeodomain-containing proteins have been intensively investigated as key molecules in the developmental toolkit. They are highly diversified and participate in a wide variety of developmental processes in metazoans. In particular, those in Cnidarians that are shared by the common ancestors of deuterostomes and protostomes are important to understand body plan evolution of bilaterians (Ferrier and Holland 2001; Chourrout et al. 2006; Ferrier 2016; DuBuc et al. 2018). While many orthologous genes of known homeodomain-containing proteins, including Hox and ParaHox genes, have been identified in cnidarians, cnidarian-specific specializations, such as loss of some homeodomain protein genes and fragmentation of the Hox cluster have been reported (Kamm et al. 2006; Steele et al. 2011; Chapman et al. 2010; Leclère et al. 2019). To understand the evolutionary trajectory of homeobox protein genes in the *Hydra* lineage, we classified them into ANTP (HOXL and NKL), PRD, LIM, POU, PROS, SINE, TALE, CERS, or ZF by bi-directional BLAST searches against sequences of homeodomains in other animals, using HomeoDB (Zhong and Holland 2011) (Table 6, Table S5) and phylogenetic analysis for ANPT- and PRD-class genes (Figs. S2 and S3), referring to the Hox genes previously identified in other cnidarians (Schummer et al. 1992; Chourrout et al. 2006; Leclère et al. 2019; Khalturin et al. 2019).

**Table 6.** Number of genes for the subclass of homeodomain-containing proteins in cnidarians.

|  | Medusozoa |  |  |  |  | Anthozoa |  |
| --- | --- | --- | --- | --- | --- | --- | --- |
|  | Hydrozoa |  |  |  |  |  |  |
| Class | *Hv* | *Hm* | *Ch* | *Mv* | *Aa* | *Nv* | *Ad* |
| ANTP-HOXL | 5 | 7 | 6 | 13 | 13 | 17 | 9 |
| ANTP-NKL | 9 | 8 | 20 | 21 | 20 | 65 | 33 |
| PRD | 21 | 16 | 18 | 25 | 28 | 43 | 31 |
| LIM | 4 | 4 | 5 | 5 | 5 | 5 | 4 |
| TALE | 4 | 3 | 5 | 4 | 10 | 6 | 5 |
| SINE | 2 | 2 | 4 | 5 | 4 | 6 | 4 |
| POU | 2 | 2 | 3 | 4 | 4 | 6 | 4 |
| CERS | 1 | 2 | 1 | 1 | 1 | 1 | 1 |
| CUT | 0 | 0 | 0 | 0 | 0 | 1 | 2 |
| HNF | 0 | 0 | 0 | 1 | 0 | 1 | 1 |
| PROS | 0 | 0 | 0 | 0 | 0 | 0 | 0 |
| ZF | 0 | 0 | 0 | 0 | 0 | 0 | 0 |
| Total | 48 | 44 | 62 | 79 | 85 | 151 | 94 |

*Hv*: *Hydra viridissima* A99, *Hm*: *Hydra magnipapillata*, *Ch*: *Clytia hemisphaerica*, *Mv*: *Morbakka virulenta*, *Aa*: *Aurelia aurita*, *Nv*, *Nematostella vectensis*, *Ad*: *Acropora digitifera*

In the *H. viridissima* genome, we identified 48 homeodomain-containing genes in the genome, 5 ANTP-HOXL, 9 ANTP-NKL, 21 PRD, 4 LIM, 4 TALE, 2 SINE, 2 POU, and 1 CERS; however, we failed to find CUT, HNF, PROS and ZF classes. This tendency toward gene loss was shared by the two other hydrozoans, *H. magnipapillata* and *Clytia hemishpaerica* (Table 6). Among cnidarians, anthozoan genomes (*Nematostella* and *Acropora*) apparently contain the most homeodomain-containing genes, while scyphozoans (*Aurelia*) and cubozoans (*Morbakka*) have intermediate numbers, and hydrozoan genomes contain the fewest. CUT class genes were not found in medusozoan genomes and all cnidarian genomes lacked PROS and ZF class genes altogether. In addition, NKL genes were less abundant in *Hydra* and HOXL genes less abundant in hydrozoans generally, than in other cnidarians.

*H.viridissima* and *H. magnipapillata* possess the same ANTP genes (Fig. 4, Table 7, Table S5), suggesting a reason for the same body plan in these *Hydra* species, although the body size of *H. viridissima* is smaller. As previously reported (Leclère et al. 2019; Khalturin et al. 2019; Gauchat et al. 2000; Quiquand et al. 2009), ParaHox genes *Gsh* and *Meox* are present in *Hydra*, whereas *Xlox* and *Cdx* are missing, unlike other medusozoans (Table 7, Fig. 4). On the other hand, Hox gene composition is quite similar among medusozoans. They have *Hox1and Hox9-14*, but lack *Hox2, Evx, Gbx, Mnx*, unlike anthozoans. Medusozoans have lost many NKL genes, *Nedx, Hlx, Mnx, Msx*, and *Lbx* compared to anthozoans. In addition, *Dbx, Hlx, Nk3, Nk6* and *Nk7* were not found in hydrozoans, nor were *Nk5, Exm* or *Msxlx* in *Hydra* (Table 7, Fig. 4). In addition, some degree of synteny conservation of HOXL genes and NKL genes is found in Anthozoa, but not in Medusozoa (Fig. 4), suggesting complete fragmentation of the homeobox gene cluster in the common ancestor of medusozoans. *Nematostella* expresses *Gbx, Hlx, Nk3* and *Nk6* in the pharyngeal or mesenteric region (*Gbx* in pharyngeal endoderm (Matus et al. 2006), *Hlx* and *Nk6* in pharyngeal ectoderm, and *Nk3* in nutrient-storing somatic gonads in mesentery (Steinmetz et al. 2017)). Anthozoans have a pharynx and a mesentery that structurally supports the pharynx and serves the site of gamete production in the gastrovascular cavity, while these tissues are not found in *Hydra*. Loss of these genes reflects the simplification of body structure in the *Hydra* lineage.

**Table. 7.**
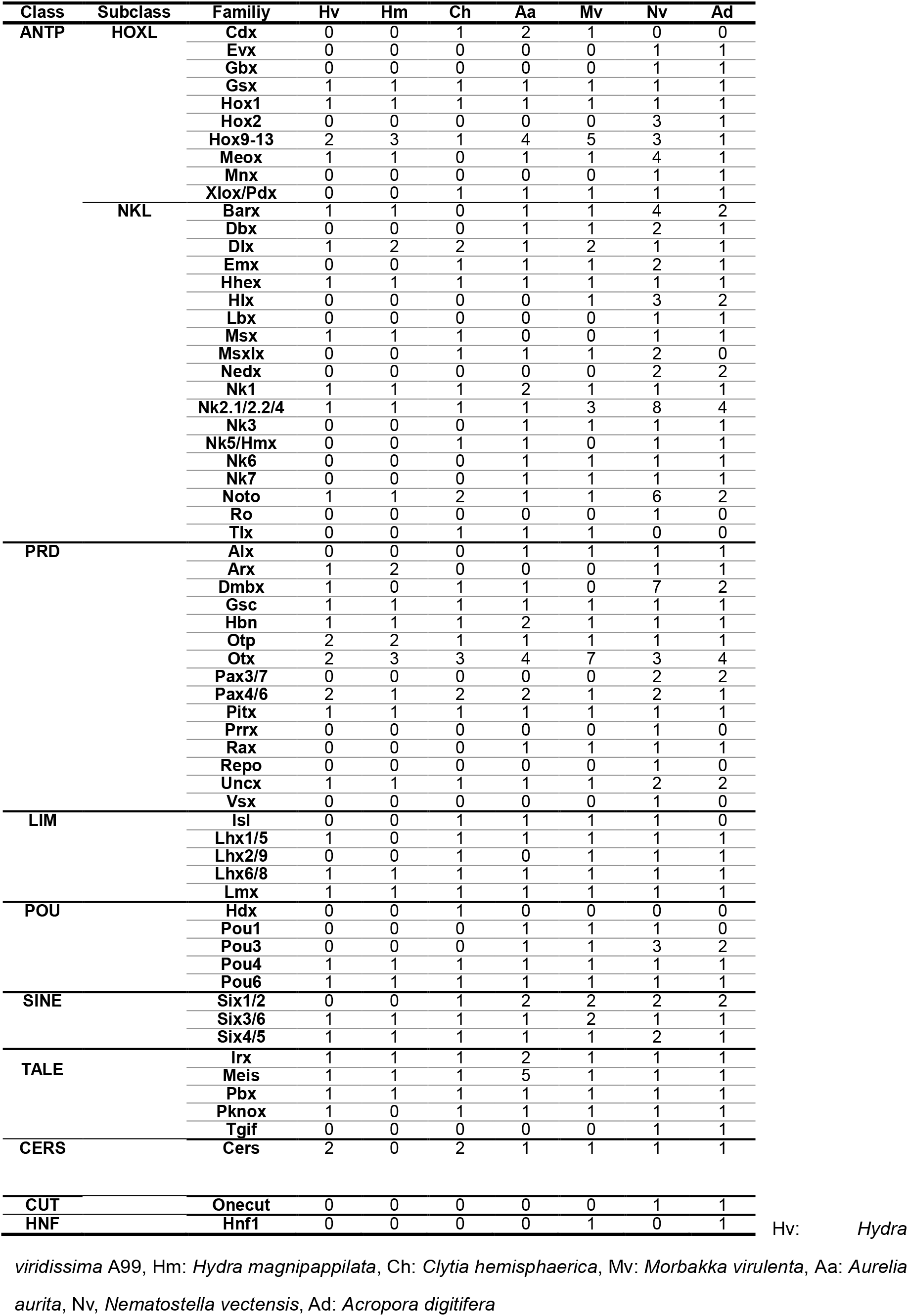
Number of homeodomain-containing genes in the *Hydra viridissima* genome.

**Figure 4.**
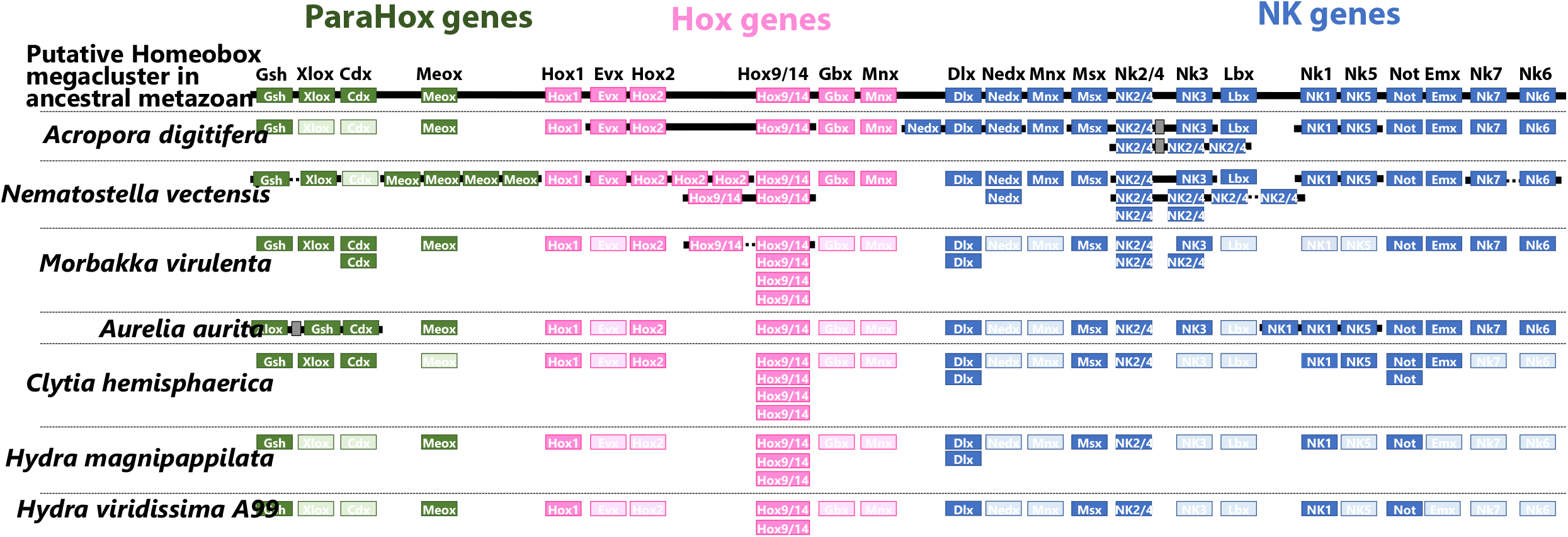
ParaHox, Hox and NK genes in cnidarians. Putative Homeobox megacluster in the last common ancestor of cnidarians and bilaterians (top) and homeobox genes and their cluster structures in present cnidarians are represented. ParaHox genes (green boxes); Hox genes (pink boxes); NK genes (blue boxes). Empty boxes indicate lost genes. Horizontal lines (black) indicate chromosome fragments.

## CONCLUSION

*H. viridissima* belongs to an early branching group of *Hydra*, which seems to have maintained the ancestral state of the genome than does more derived *H. magnipapillata.* On the other hand, the repertoire of transcriptional factor genes including homeodomain-containing genes in *H. viridissima* is quite similar to that in *H. magnipapillata*, reflecting the common body plan in these species. Comparative analysis of transcription factor genes among cnidarians indicates gradual simplification of the ANTP gene repertoire in the *Hydra* lineage, which is likely to reflect the simple body structure of *Hydra*. In addition, we found diverse innate immunity genes in *H. viridissima* genome that are also observed in coral, indicating the common feature involved in algal symbiosis. The *H. viridissima* genome presented here provides a *Hydra* genome comparable in quality to those of other cnidarians, including medusozoans and anthozoans, which will hopefully facilitate further studies of cnidarian genes, genomes, and genetics to understand basal metazoan evolution and strategies to support algal symbiosis in cnidarians.

## ACKNOWLEDGEMENTS

This research was supported by funding provided by Okinawa Institute of Science and Technology (OIST) to the Marine Genomics Unit (NS) and by the Okayama University Dispatch Project for Female Faculty members (MH). MH was supported by Japanese Society for Promotion of Sciences Funding (15K07173, 18K06364). We thank Dr. Steven D. Aird for editing the manuscript, Ms. Kanako Hisata (OIST) for creating the genome browser, Dr. Chuya Shinzato (OIST, Tokyo University) for providing valuable advice and help with genome sequencing and assembly, and Prof. Thomas C. G. Bosch (Kiel University) for offering valuable discussion. We also thank the DNA Sequencing Section and the IT Section of OIST for excellent technical support. Computations for this work were partially performed on the NIG supercomputer at the ROIS National Institute of Genetics.

